# IPSE, a parasite-derived host immunomodulatory protein, is a promising therapeutic for hemorrhagic cystitis

**DOI:** 10.1101/400424

**Authors:** Rebecca S. Zee, Evaristus C. Mbanefo, Loc H. Le, Luke F. Pennington, Justin Odegaard, Theodore S. Jardetzky, Abdulaziz Alouffi, Jude Akinwale, Franco H. Falcone, Michael H. Hsieh

## Abstract

Chemotherapy-induced hemorrhagic cystitis is characterized by bladder pain and voiding dysfunction caused by hemorrhage and inflammation. Of currently available therapies, prophylactic 2-mercaptoethanesulfonic acid (MESNA) has limited efficacy and cannot treat pre-existing lesions. Therefore, novel therapeutic options to treat hemorrhagic cystitis are needed. We previously reported that systemic administration of the *Schistosomiasis haematobium*-derived protein H-IPSE^H06^ (IL-4-inducing <u>p</u>rinciple from *<u>S</u>chistosoma mansoni* <u>e</u>ggs), is superior to 3 doses of MESNA in alleviating hemorrhagic cystitis. Based on prior reports by others on *S. mansoni* IPSE and additional work by our group, we reasoned that H-IPSE^H06^ mediates its effects on hemorrhagic cystitis by binding IgE on basophils and inducing IL-4 expression, promoting urothelial proliferation, and translocating to the nucleus to modulate expression of genes implicated in relieving bladder dysfunction. We speculated that local bladder injection of the *S. haematobium* IPSE ortholog IPSE^H03^, hereafter called H-IPSE^H03^, might be more efficacious in preventing hemorrhagic cystitis compared to systemic administration of IPSE^H06^. We demonstrate herein that H-IPSE^H03^ is a promising therapeutic for the treatment of voiding dysfunction and bladder pain in hemorrhagic cystitis. Namely, it attenuates ifosfamide-induced increases in bladder wet weight in an IL-4-dependent fashion. H-IPSE^H03^ relieves hemorrhagic cystitis-associated allodynia. Finally, H-IPSE^H03^ drives increased urothelial cell proliferation. This indicates that IPSE induces bladder healing mechanisms, which suggests that it may be a novel non-opioid analgesic to treat bladder pain syndromes.

## Introduction

Ifosfamide and other alkylating chemotherapy agents are used in a wide variety of malignancies including leukemias, soft tissue sarcomas, and testis cancer. The liver metabolizes ifosfamide into acrolein, which is excreted in the urine and has a deleterious effect on the urothelium. Hemorrhagic cystitis is characterized by bladder edema, hemorrhage, urothelial denudation, and infiltration of inflammatory cells. This condition affects up to 40% of ifosfamide-exposed patients, resulting in hematuria, dysuria, bladder spasms, and urinary frequency (9). Hemorrhagic cystitis is a challenging condition to manage, and often requires hospitalization and invasive treatments (16).

Accordingly, strategies to attenuate ifosfamide-induced hemorrhagic cystitis, such as administration of 2-mercaptoethanesulfonic acid (MESNA), bladder irrigation, or hyperhydration often achieve suboptimal protection for patients (16). Despite use of existing therapies, a majority of patients have symptomatic and/or histologic evidence of hemorrhagic cystitis (14). As an alternative to current management approaches, Macedo *et al*. reported that administration of recombinant interleukin-4 (IL-4) attenuated the effects of ifosfamide in a mouse model of hemorrhagic cystitis (15). The importance of IL-4 in this model was demonstrated by administration of anti-IL4 antibody to ifosfamide-exposed, wild type mice and administration of ifosfamide to IL-4-deficient mice, both of which resulted in worsened hemorrhagic cystitis (20). Interestingly, ifosfamide administration increased endogenous production of IL-4, suggesting the existence of intrinsic regulatory mechanisms to control inflammation in response to ifosfamide (15). However, systemic administration of IL-4 to treat hemorrhagic cystitis may not be a realistic option due to pleiotropic effects and a short *in vivo* half-life of this cytokine (18). Therefore, alternative strategies to increase expression of IL-4 would be needed in order to leverage this cytokine for therapeutic treatment of hemorrhagic cystitis.

One alternative may be the interleukin-4 inducing principle from *Schistosoma mansoni* eggs (IPSE), the most abundant protein secreted by *S. mansoni* eggs. IPSE attenuates inflammation via multiple mechanisms, including binding immunoglobulins to stimulate IL-4 release, sequestering chemokines, and translocating to the nucleus to modulate transcription (17, 19). We have previously reported that similar to the *S. mansoni* ortholog of IPSE, M-IPSE, several *S. haematobium* orthologs, referred hereafter as H-IPSE, bind to IgE on mast cells and basophils and upregulate the expression of IL-4 (19). We identified two main clades of H-IPSE exemplified by the orthologs IPSE^H03^ and IPSE^H06^. Importantly, both IPSE^H03^ and IPSE^H06^ translocate into urothelial cell nuclei (19).

Initial animal experiments with H-IPSE focused on the effect of systemic administration of H-IPSE^H06^ by tail vein injection (17). Tail vein injection of H-IPSE^H06^ attenuates ifosfamide-induced bladder hemorrhage in an IL-4 and NLS-dependent manner. Furthermore, mice treated with H-IPSE^H06^ prior to ifosfamide exposure demonstrated fewer spontaneous pain behaviors and had a higher threshold for evoked pain responses. We speculated that direct injection of IPSE into the bladder wall would have multiple advantages over intravenous injection, including avoidance of side effects caused by systemic administration (although none have been identified to date), and potentially decreased dosage to achieve a therapeutic effect. The aim of this work was to determine whether direct bladder wall injection of H-IPSE^H03^ attenuates bladder inflammation, voiding dysfunction and pain in a mouse model of hemorrhagic cystitis.

## Materials and Methods

### Mice

Six to 8-week-old female C57BL/6 mice (Charles River Laboratories, Wilmington, MA) were housed in cages with free access to water and standard chow and 12 hour light-dark cycles. Mice were acclimated for at least 7 days prior to experimentation. The animal protocol was approved by the Institutional Animal Care and Use Committee at the Biomedical Research Institute (Rockville, MD). Our institutional animal care and use committee guidelines follow the U.S. Public Health Service Policy on Human Care and Use of Laboratory Animals.

### Bladder wall injections

Mice were anesthetized with 2% continuous isoflurane on a heating pad. Procedures were performed using sterile technique. For pain control, 0.1 mg/kg buprenorphine and 0.1 mg/kg bupivacaine were injected subcutaneously. A midline laparotomy was performed sharply and the bladder delivered through the incision. Mice were divided into 3 groups receiving sham, control or IPSE. A 30-gauge needle was used to inject a 1:1 v/v mixture of Low Growth Factor Matrigel (Corning, Corning, New York) and PBS containing 25 μg mouse albumin (control) or 25 µg H-IPSE (IPSE) (Figure 1). Sham mice received a midline laparotomy only. Incisions were closed in 2 layers using 5-0 Vicryl on the abdominal wall and 5-0 silk to close skin. Bacitracin was applied to the incision. The mice were recovered on a heating pad. Twenty-four hours later mice were injected with 400 mg/kg ifosfamide (Sigma-aldrich, St. Louis, MO). Mice who received anti-IL4 antibody (inVivoMab 11B11, BioXcell, West Lebanon, NH) received 10 ng by intraperitoneal (IP) injection 30 24 minutes before ifosfamide. Control mice received IP injections of phosphate-buffered saline (PBS). At 12 hours, mice were euthanized, bladders were removed and weighed. Bladders were then subjected to additional analysis detailed below.

**Figure 1:**
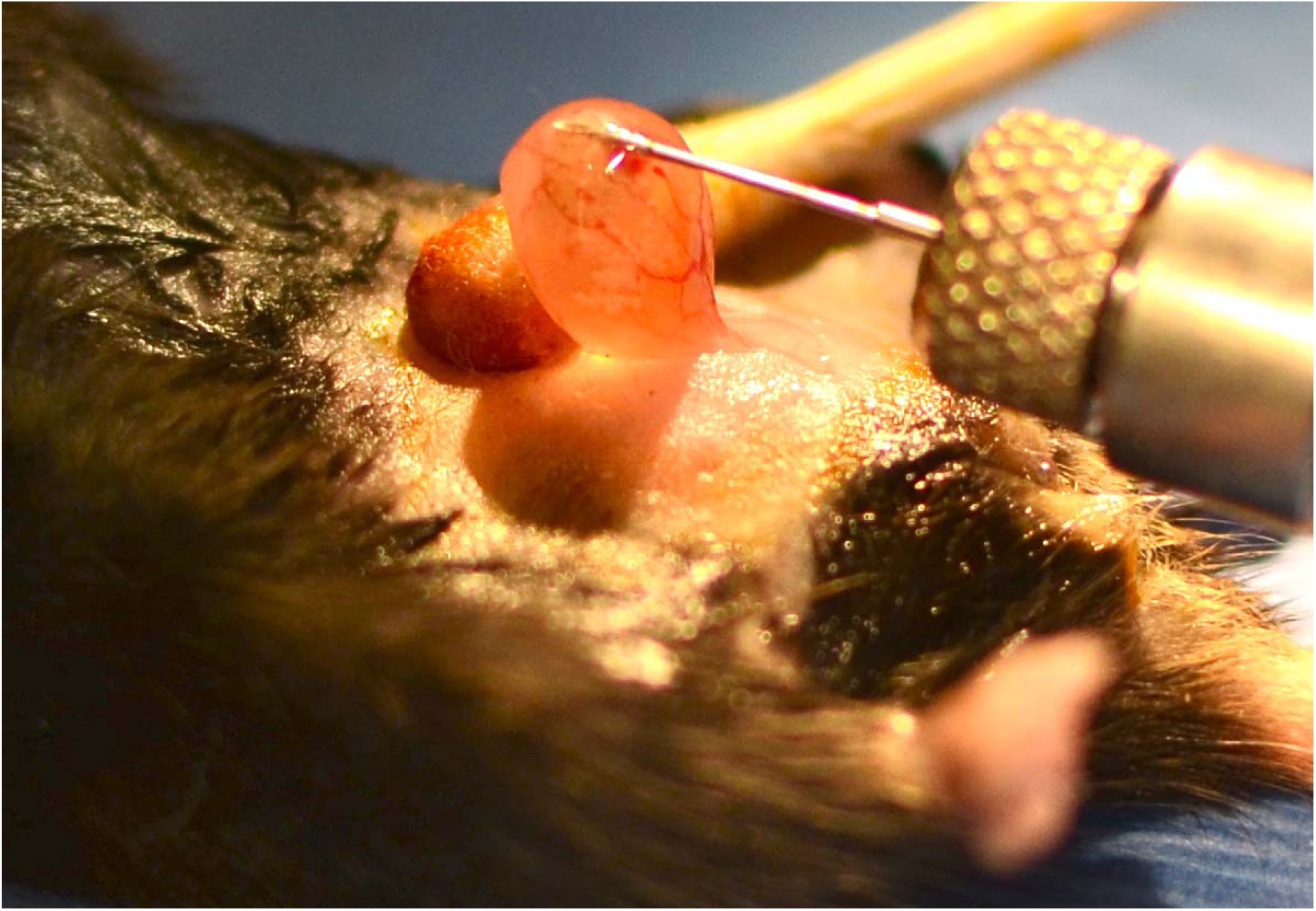
Bladder wall injection technique. Mice were anesthetized with inhaled isoflurane. Next their abdomens were depilated and cleaned, injected with local anesthetic, and a midline laparotomy was performed. The bladder was exteriorized, stabilized with a cotton applicator, and its wall injected with a 30 gauge needle.

### Tail vein injections

Mice were anesthetized with 2% continuous isoflurane on a heating pad. A 30-gauge needle was used to inject PBS containing 25 µg mouse albumin or 25 µg H-IPSE^H03^ (IPSE) in PBS. The mice were recovered on a heating pad. Twenty-four hours later mice were injected with 400 mg/kg ifosfamide (Sigma-aldrich, St. Louis, MO). Mice who received anti-IL4 antibody (inVivoMab 11B11, BioXcell, West Lebanon, NH) received 10 ng by intraperitoneal (IP) injection 30 minutes before ifosfamide. Control mice received IP injections of phosphate-buffered saline (PBS). At 12 hours, mice were euthanized, bladders were removed and weighed. Bladders were then subjected to additional analysis detailed below.

### Recombinant IPSE protein

Recombinant IPSE protein was generated as previously described (1,2). One milligram of plasmid DNA was purified using a GeneElute HP endotoxin-free plasmid Maxiprep kit (Sigma-Aldrich), and incubated with 3 mg linear 25 kDa polyethylenimine (PolySciences, Warrington, PA) at 1 mg/mL. Finally, the plasmid was diluted in 10 mL sterile PBS for each transfection in 1L. Human embryonic kidney 293-6E cells (7) expressed secreted recombinant protein for 5 days in suspension culture using FreeStyle 293 Medium (Thermo Fisher Scientific, Waltham, MA, USA) (Figure 2A). Protein was purified over 10 mL Ni-NTA resin (Qiagen, Germantown, MD, USA), washed with 25 mM imidazole PBS, pH 7.4, and eluted with 300 mM imidazole PBS, pH 7.4 containing 50 mM arginine. Eluted protein was concentrated with an Amicon Ultra Centrifugal Filter Unit (EMD Millipore, Billerica, MA, USA) followed by purification with a Hiload 16/600 Superdex 200 Column (GE Healthcare, Waukesha, WI, USA). Nuclear localization mutants were generated using site-directed mutagenesis. These mutants (124-PKRRRTY-130 to 124-PKAAATY) disrupted the C-terminal NLS (NLS; H-IPSE^H03NLS^) (2). To decrease the risk of pyrogen contamination, FPLC machines and Hiload columns were cleaned with 0.5 M NaoH for greater than 2 hours of continuous flow and then washed with PBS, pH 7.4.

**Figure 2:**
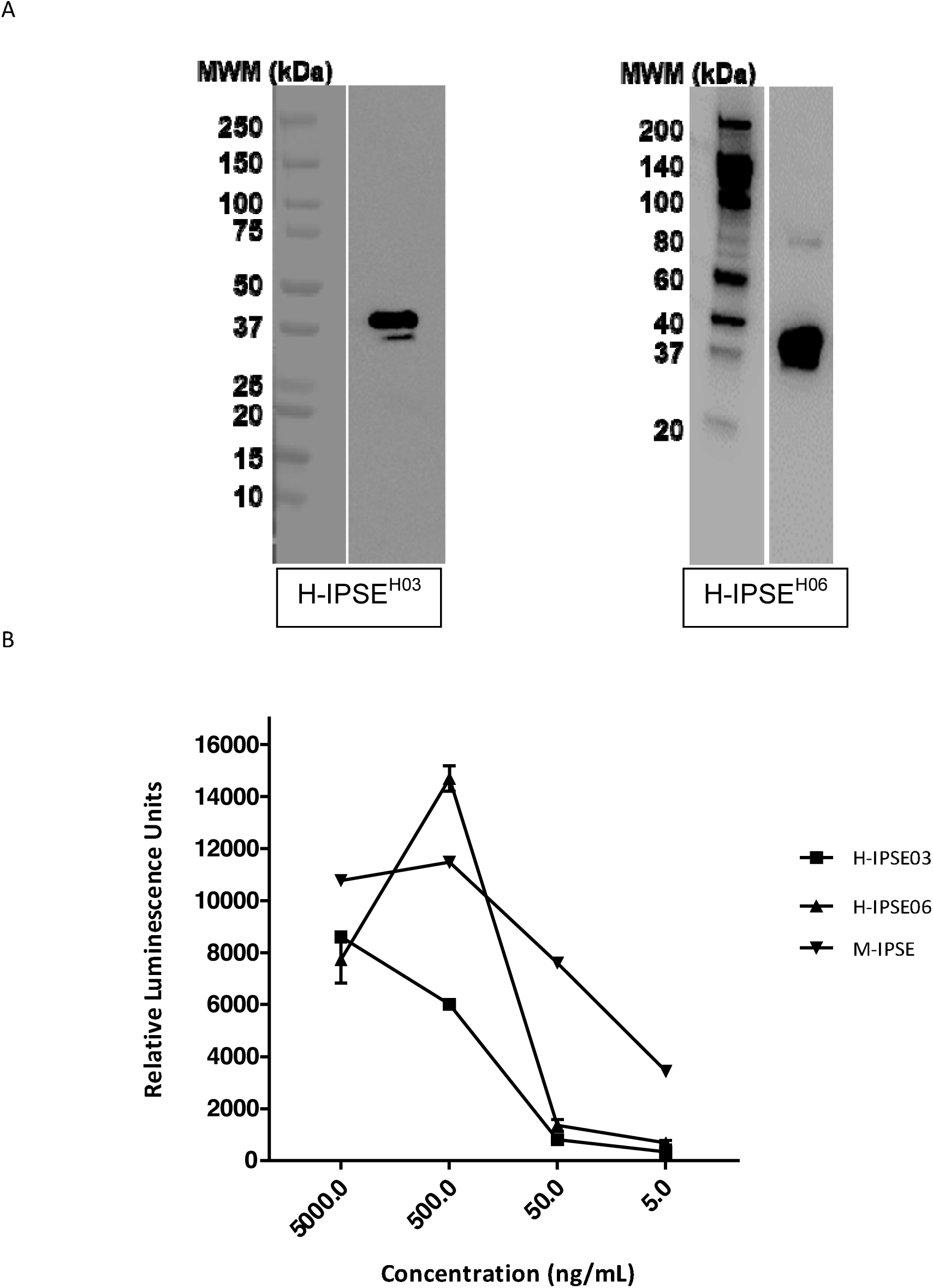
(A) Western blots of purified H-IPSE^H03^ and H-IPSE^H06^ proteins demonstrate a prominent band with a molecular weight of approximately 38-40 kDa. (B) Basophil NF-AT activation in response to M-IPSE, H-IPSE^H03^ and H-IPSE^H06^. Incubation of IPSE with IgE-bearing basophils demonstrates that H-IPSE orthologs induce NF-AT reporter gene expression comparable to M-IPSE.

### SDS-PAGE and Western blotting

Purified protein was separated on 4-20% gradient gels by SDS-PAGE in 15 µL aliquots (Mini-Protean TGX Precast Gels, Biorad). Separated proteins were then transferred to a 0.2 µM nitrocellulose membranes. Membranes were incubated in blocking buffer for 1 hour (5% [wt/vol] dried skim milk, 0.01% [vol/vol] Tween 20, and Tris-buffered saline [TBS]) on a shaker at room temp. Primary antibody was mouse anti-His (GE-Healthcare) diluted at 1:500 and incubated overnight at 4°C followed by washing in TBS containing 1% Tween 20 for 5 min x 3. Membranes were then incubated with secondary antibody--HRP-conjugated anti-mouse IgG (Sigma-Aldrich) -- for 1 hour at room temperature followed by 3 additional washes. Imaging was performed with chemiluminesence-luminol reagent (3 µL of 30% H_2_O_2_, 0.1 Tris-HCl [pH 8.0], 2.5 mM luminol, and 400 µM coumaric acid) on a Fuji LAS4000 imager.

### Basophil activation with recombinant M-IPSE, H-IPSE^H03^ and H-IPSE^H06^

Basophil activation with recombinant M-IPSE, H-IPSE^H03^ and H-IPSE^H06^ Basophil activation was quantified as previously described (23). RS-ATL8 cells were cultured in 10 mL MEM (GIBCO, USA), supplemented with 5% vv v/v heat-inactivated FCS (GIBCO, USA), 100 U/mL penicillin, and 100 µg/mL streptomycin (Sigma, UK) and 2 mM L-glutamine (Sigma, UK). Medium was changed every 2-3 days. Cells were grown in 75 cm^2^ flasks at 37°C in a humidified atmosphere with 5% carbon dioxide. 1 mg/mL G418 (Fisher ThermoScientific, UK) and 600 µg/mL hygromycin B (Invitrogen, Paisley, UK) were used to maintain expression of human FcεRI genes and NFAT-luciferase, respectively. Prior to testing, cells were incubated overnight with M-IPSE, H-IPSE^H03^ or H-IPSE^H06^ at concentrations ranging from 5 to 5000 ng/mL. Luciferase assays were performed with ONE-Glo Luciferase Assay System (Promega, UK), following the manufacturer’s instructions. The luciferase substrate was added and chemiluminescence was measured using an Infinite M200 microplate reader (Tecan, MÛnnedorf, Switzerland) within 30 minutes.

### Pain assessment

Visceral pain scores were assigned as previously described (10). The observer was blinded to mouse treatment assignments prior to assessments. Mice were placed in clean cages and acclimated for 30 min. For spontaneous pain scoring, mice were observed for 60 seconds and given a cumulative spontaneous pain score based on the following: (0) – normal; (1) – piloerection; (2) – labored breathing; (3) – ptosis; (4) – licking of abdomen (not grooming); (5) – rounded back. The maximum possible visceral pain score is 15. Pain scores were collected at baseline (prior to bladder wall injection), and 10 hours after ifosfamide was administered.

### Von Frey filament testing

Evoked pain scores were collected in a blinded fashion to assess for referred hyperalgesia. We adopted the up-down approach as previously described (6, 13). An electronic Von Frey filament (BioSeb, Pinellas Park, Florida) was applied to the right hind footpad of the mouse for 5 seconds until the mouse displayed rapid withdrawal of the paw, jumping, or licking of the paw. The 50% withdrawal threshold was then calculated from an average of 3 measurements. Results are tabulated as the difference between baseline and post-ifosfamide values.

### Voided Spot on Paper Assay

Voided spot on paper assays were performed as previously described (1, 8, 11, 25). Mice were placed in individual cages 2 hours after ifosfamide or PBS administration. Whatman paper was cut to the dimensions of the cage floor. The paper was covered with wire mesh to prevent mice from tearing or ripping the paper. Food was provided *ad libitum* in the form of regular chow. Water was not provided to prevent fluid dripping onto the paper and causing data loss or artifact. Mice were placed under quiet conditions for 4 hours. They were then returned to normal housing conditions after completion of the experiment. The pieces of Whatman paper were converted to .tiff images using UV transillumination (Bio-Rad, Hercules, CA). Image analysis was performed with ImageJ Fiji (https://fiji.sc/). Corner voiding was assessed by assigning 5% of the total paper area to each corner. Central voiding was assessed by assigning 40% of the total area to the center of the filter paper.

### *In vitro* proliferation assays

MB49 cells were counted and plated with equal numbers of cells in each well. H-IPSE^H03^ or H-IPSE^H03NLS^ were added to the cell media at the following concentrations: 0.0655 pmol (1 ng/ml), 0.655 pmol (10 ng/ml), 6.55 pmol (100 ng/ml), 65.5 pmol (1000 ng/ml), or PBS for control. 5-(and 6)-Carboxyfluorescein diacetate succinimidyl ester (CFSE) assays were then performed according to manufacturer’s instructions (Thermofisher Scientific, Waltham, MA). One mL of a single cell suspension for each experimental condition was then acquired on a BD FACSCanto II machine (BD Biosciences, San Jose, CA). Flow cytometric analysis was performed using FlowJo software (Ashland, OR).

### Statistical analysis

One-way ANOVA or Student’s t-test were utilized as appropriate. *Post hoc* testing was performed with Bonferroni test. A p-value of less than 0.05 was considered statistically significant.

## Results

### Recombinant H-IPSE^H03^ and H-IPSE^H06^ proteins activate IgE-bearing basophils *invitro*

We previously demonstrated that M-IPSE activates basophils *in vitro* through NF-AT (23). This pathway is implicated in basophil and mast cell expression of IL-4, which we have observed *in vivo* in mice administered H-IPSE^H06^ (17). Moreover, we have also noted that ifosfamide-challenged mice given H-IPSE^H06^ are protected from several pathogenic aspects of hemorrhagic cystitis in an IL-4-dependent fashion (17). Thus, we sought to demonstrate that H-IPSE^H03^ and H-IPSE^H06^, which are both *S. haematobium* orthologs of M-IPSE, also stimulate IL-4-associated reporter gene expression *in vitro*. We first purified recombinant H-IPSE^H03^ and H-IPSE^H06^ from transfected HEK293-6A cells. Western blots identified a band with a molecular weight of 38-40 kDa which corresponds to the homodimeric H-IPSE structure (Figure 2A). Recombinant H-IPSE^H03^ and H-IPSE^H06^ protein was then incubated with IgE-loaded basophils. This resulted in NF-AT activation, which is associated with IL-4 secretion in basophils (Figure 2B). Having confirmed that H-IPSE^H03^ triggers IL-4-associated pathways in cultured basophils, we next sought to determine the therapeutic efficacy of H-IPSE^H03^ in the mouse model of ifosfamide-induced hemorrhagic cystitis.

### H-IPSE^H03^ dampens chemotherapy-induced increases in bladder wet weight

We assessed for an increase in bladder wet weight caused by hemorrhage, edema and cellular infiltration following ifosfamide injection. Ifosfamide administration caused a statistically significant increase in bladder wet weight compared to controls (Figure 2A, 2B; n=8, p<0.001) H-IPSE^H03^ bladder wall injection significantly reversed the increase in bladder wet weight caused by ifosfamide in bladder wall injected mice but not mice that received tail vein injection (p=0.02 and N.S., respectively). The beneficial effect of IPSE on bladder wet weight was reversed by anti-IL4 antibody (p<0.001). However, IPSE^H03NLS^ also ameliorated ifosfamide-induced increases in bladder wet weight, regardless of mode of administration, suggesting that the therapeutic effect of IPSE on bladder wet weight is mediated by IL-4, but not dependent on IPSE translocation into the nucleus.

Tail vein injection results were distinct from bladder wall injection in two ways. After tail vein H-IPSE^H03^ injection bladder wet weights decreased but remained significantly higher than non-ifosfamide-exposed controls (Figure 2B; p=0.03). Furthermore, administration of H-IPSE^NLS^ to ifosfamide-treated mice demonstrated a downward trend in bladder wet weight that was not significant compared to mice given only ifosfamide.

### H-IPSE^H03^ abrogates evoked pain responses in chemotherapy-treated mice

We next sought to determine whether H-IPSE^H03^ administration had an effect on ifosfamide-induced bladder pain. We first measured referred hyperalgesia using von Frey filament testing. Mice injected with ifosfamide had greater evoked pain responses than those of control mice (Figure 3). H-IPSE^H03^ bladder wall injection increased the withdrawal threshold, i.e., reversed allodynia caused by ifosfamide injection (p<0.05). When neutralizing anti-IL-4 antibody was co-administered with H-IPSE^H03^, the protective effect of H-IPSE^H03^ was attenuated (p<0.05). Likewise, injection of H-IPSE^H03NLS^, which cannot translocate to the nucleus, also featured a decreased analgesic effect compared to IPSEH03 (p<0.05). H-IPSE^H03^ had no effect on referred hyperalgesia when administered via tail vein injection (data not shown)

**Figure 3:**
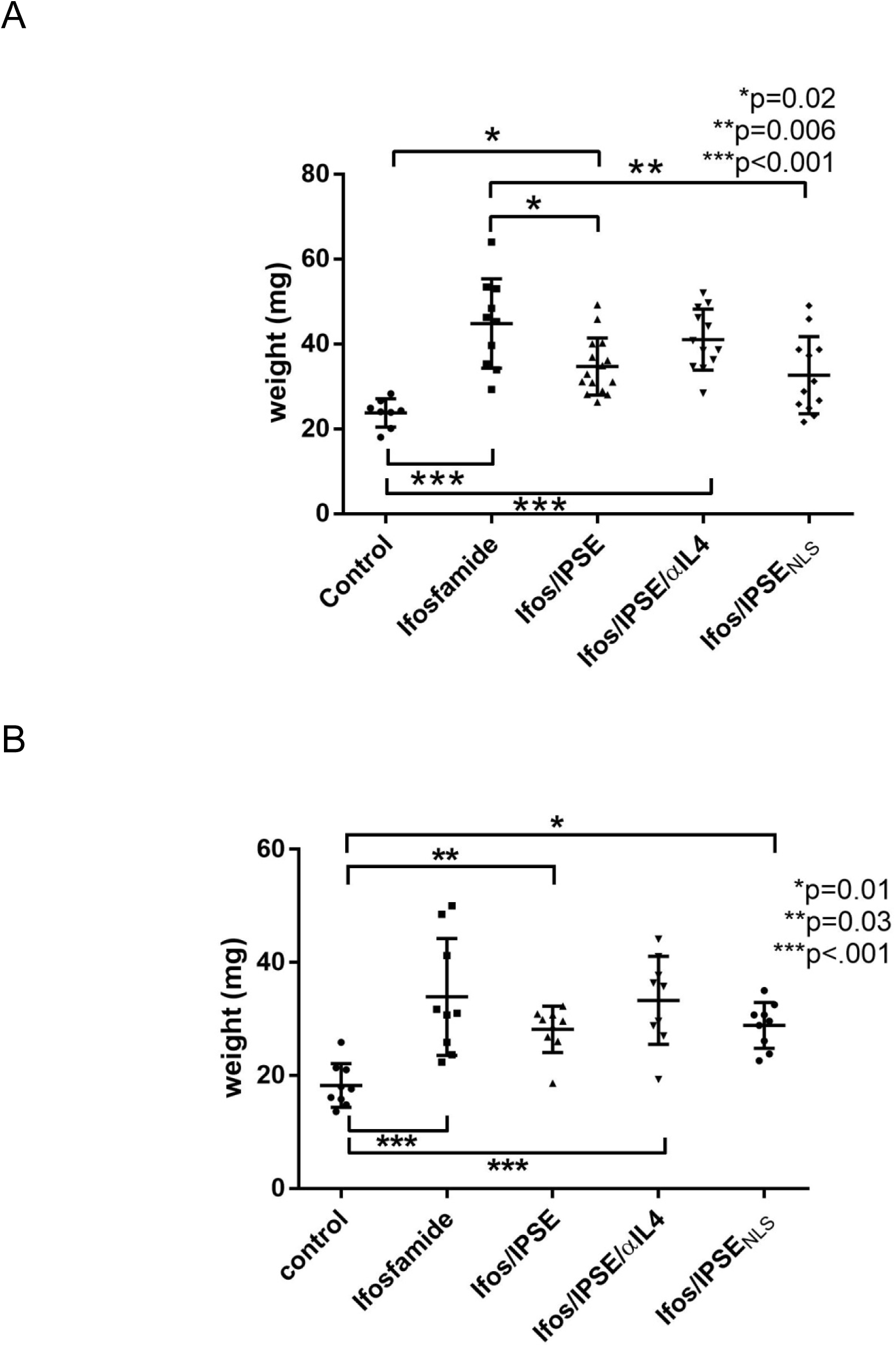
Effect of H-IPSE^H03^ on bladder wet weights. Mice received a bladder wall injection (A) or a tail vein injection (B) with or without H-IPSE^H03^ or a nuclear localization sequence mutant of H-IPSE^H03^ (IPSE_NLS_). Twenty-four hours later, mice were injected with PBS (control) or ifosfamide (“ifosfamide” or “ifos”). Some mice received neutralizing anti-IL4 antibody (αIL4) 30 minutes prior to ifosfamide. Bladders were collected and weighed 12 hours following ifosfamide injection to assess for edema and hemorrhage. **A**. Bladder wall injection of H-IPSE^H03^ significantly decreases ifosfamide-induced increase in bladder wet weight in an IL-4-but not nuclear translocation-dependent fashion. **B.** Tail vein injection of H-IPSE^H03^ non-significantly decreases the ifosfamide-induced increase in BWW in an IL-4 but not NLS-dependent fashion. Plotted data are pooled from 3 experiments. Error bars represent standard deviations. hyperalgesia). H-IPSE^H03^ bladder wall injection alleviates allodynia (referred hyperalgesia) associated with hemorrhagic cystitis-associated pain in an IL-4 and NLS-dependent manner. Plotted data are pooled from 3 experiments. (*p=0.04, **p=0.02, ***p=0.01; Bars represent means and one standard deviation)

### H-IPSE^H03^ does not significantly affect abnormal ifosfamide-induced voiding patterns in mice

Mice are prey animals and preferentially void in the corner of their enclosures as a predator avoidance strategy. We assessed for voiding dysfunction caused by ifosfamide based on the percentage of overall voids in the corners of cages. When mice received ifosfamide, the percentage of corner voids was significantly decreased (Figure 4A). H-IPSE^H03^ increased the frequency of corner voiding in the presence of ifosfamide, but this was not a statistically significant finding. Administration of α-IL4 antibody reversed the effect of H-IPSE^H03^ on corner voiding and was not significantly different from ifosfamide treatment (p=0.07 vs. control).

**Figure 4:**
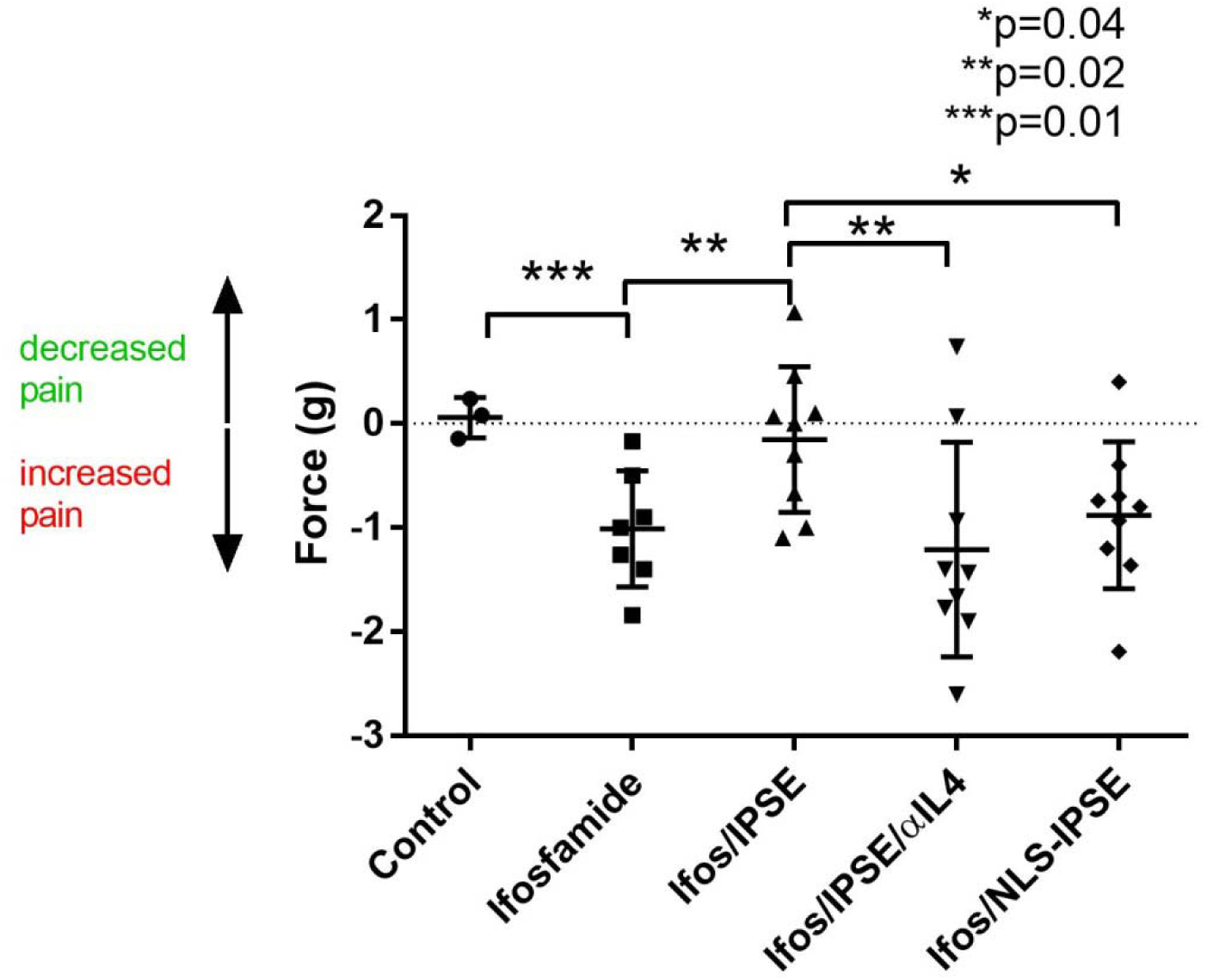
The effect of IPSE bladder wall 23 injections on evoked pain responses (referred hyperalgesia). H-IPSE^H03^ bladder wall injection alleviates allodynia (referred hyperalgesia) associated with hemorrhagic cystitis-associated pain in an IL-4 and NLS dependent manner. Plotted data are pooled from 3 experiments. (*p=0.04, **p=0.02, ***p=0.01; Bars represent means and one standard deviation)

Ifosfamide administration non-significantly increased the percentage of voids in the central area of cages (Figure 4B). H-IPSE^H03^, α-IL4 antibody or H-IPSE^H03NLS^ did not have a significant effect on central voiding. H-IPSE^H03NLS^ -treated mice were not significantly different from ifosfamide-treated or control mice. Tail vein injection of H-IPSE^H03^ did not significantly improve or alter voiding patterns in ifosfamide-treated mice (Data not shown).

### H-IPSE^H03^ promotes proliferation of urothelial cells *in vitro*

Given the beneficial effects of H-IPSE^H03^ on ifosfamide-induced bladder wet weight increases and pain, as well as prior data indicating a direct effect of H-IPSE^H03^ on urothelial cells (17), we assessed the effect of H-IPSE^H03^ on urothelial cell IPSE^H03^ significantly increased cell proliferation over two successive daughter cell proliferation by co-incubating H-IPSE^H03^ with the MB49 (mouse urothelial) cell line. H-IPSE^H03^ generations compared to controls (Figure 5A; *p<0.05, **p<0.01, ***p<0.0001; n=8). This held true across a range of H-IPSE^H03^ concentrations. In contrast, co-incubation of cells with H-IPSE^H03NLS^ did not cause increased proliferation over that of controls (Figure 5B).

**Figure 5:**
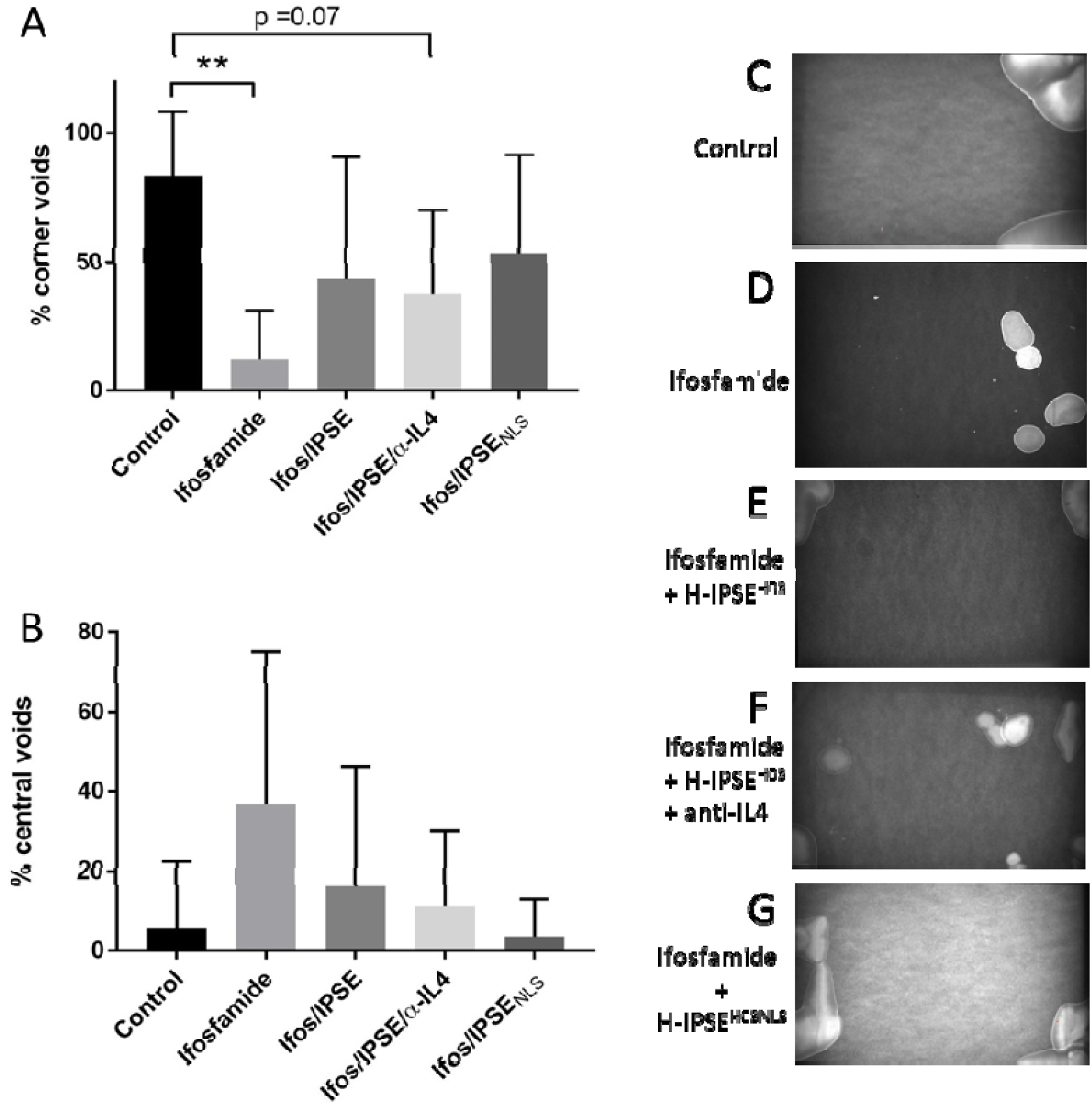
Voiding dysfunction caused by ifosfamide was non-significantly alleviated bybladder wall injections of H-IPSE^H03^. Ifosfamide significantly decreased corner voiding(A & D). H-IPSE^H03^ non-significantly restored the percentage of corner voids inifosfamide-treated mice (E). Administration of neutralizing α-IL4 antibody may reversethe non-significant protective effect of H-IPSE^H03^ (F). Central voiding tended to increase following ifosfamide administration (B). The additionof H-IPSE^H03^ (E), neutralizing α-IL4 antibody (F) or H-IPSE^H03NLS^ (G) did not significantlychange the percentage of voiding observed in the center of the cage. Plotted data arepooled from 3 experiments. **p<0.01; error bars represent standard deviations.

**Figure 6:**
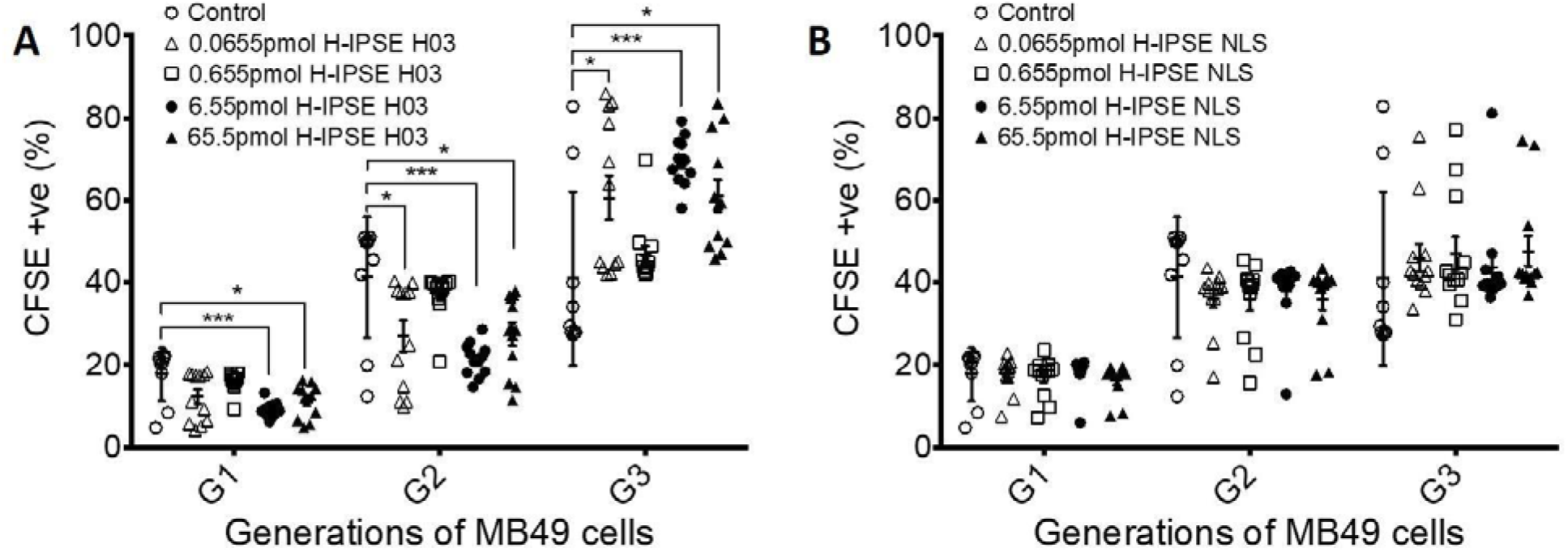
H-IPSE^H03^ co-incubation with MB49 cells induced proliferation in an NLS-dependent fashion. (A) When co-incubated with H-IPSE^H03^, the number of MB49 cellswas markedly increased versus control over 3 generations of cells. Significant increasesin proliferation were observed for both low (0.065 pmol) and high concentrations of H-IPSE^H03^ (up to 65.5 pmol). (B) MB49 cellular proliferation was not increased comparedto controls by co-incubation with H-IPSE^H03NLS^.

## Discussion

Hemorrhagic cystitis is a common sequela of alkylating chemotherapy, affecting up to 40% of patients who receive ifosfamide or cyclophosphamide (14). Once established, hemorrhagic cystitis is a challenging-to-manage entity characterized by widespread bladder inflammation and leading to hematuria, dysuria, small volume voids, urinary frequency, and bladder spasms. Currently available medical therapy, MESNA, has a narrow therapeutic window as it can only be administered immediately before and during chemotherapy. MESNA can cause hypersensitivity reactions and is ineffective in treating hemorrhagic cystitis once it has been established (2, 21, 22). Therefore, novel therapies need to be developed to fulfill this unmet need.

One source of new drugs for hemorrhagic cystitis may be derived from *Schistosoma haematobium.* Urogenital schistosomiasis is a parasitic disease in which *Schistosoma haematobium* worms lay eggs in the bladder and other pelvic organs. Deposited eggs must traverse the host bladder wall in order to be released in the urine. Although urogenital schistosomiasis itself causes a form of hemorrhagic cystitis, Hematuria can be variable or even absent (24). We reasoned that host immunomodulation by *S. haematobium* egg products allow the parasite to complete its life cycle without causing severe morbidity to its host, including hemorrhagic cystitis (10). Specifically, we postulated that *S. haematobium* eggs can accomplish this by secreting H-IPSE orthologs in order to modulate the host immune response.

In a prior study we demonstrated the clinical potential of exploiting the anti-inflammatory and analgesic properties of H-IPSE^H06^ (17). A single intravenous dose of H-IPSE^H06^ was superior to MESNA in alleviating bl adder hemorrhage in ifosfamide treated mice (17).

Clinical translation of H-IPSE^H06^, H-IPSE^H03^, and other IPSE orthologs will require large-scale recombinant protein production. Herein we show that H-IPSE^H03^ and H-IPSE^H06^ can be purified from mammalian HEK293T-6A cells. Furthermore, we demonstrate that, like M-IPSE, H-IPSE^H03^ and H-IPSE^H06^ trigger IgE-bearing basophil NF-AT activation *in vitro*, which in turn is linked to IL-4 secretion. H-IPSE^H03^ injected into the mouse bladder wall attenuates ifosfamide-induced increases in bladder wet weight in an IL-4 and NLS-dependent fashion. This suggests that H-IPSE^H03^ reduces ifosfamide-induced edema, cellular infiltration, and/or hemorrhage (pathologic processes which can increase bladder wet weight). When H-IPSE^H03^ was administered via tail vein, ifosfamide-induced increases in bladder wet weight were unaffected. Unsurprisingly, there was neither an IL-4-dependent nor nuclear translocation-dependent effect compared to controls. This is consistent with our prior report that intravenous administration of H-IPSE^H06^ did not affect ifosfamide-mediated increases in bladder wet weight (17). There are several possible explanations for these differences in effects of bladder wall versus intravenous injections. For instance, bladder wall injections themselves may cause an increase in hemorrhage, and bladder mass due to the added weight of the matrigel. This may make it more difficult to discern weight differences between groups when compared to tail vein injection. Furthermore, bladder wall injection of H-IPSE^H03^ may result in high local concentrations but low systemic levels. We have previously reported that peripheral basophils may play a role in IPSE’s therapeutic effects in hemorrhagic cystitis (17). Recruitment of circulating basophils to the site of inflammation and subsequent IL-4 release may be dependent on the action of H-IPSE outside of the bladder. Conversely, it is possible that the higher local H-IPSE^H03^ concentrations achieved by bladder wall injection may more effectively activate bladder mast cells, basophils, and other cell types critical for therapeutic effects.

Another explanation for the different phenotypes observed between H-IPSE^H03^ and H-IPSE^H06^ is that variations in the sequence, and therefore, function of IPSE proteins have evolved such that different orthologs of H-IPSE are secreted to perform different host-modulatory functions. Both orthologs of H-IPSE are homologous to M-IPSE in that they both conserve the C-terminal nuclear localization sequence as well as 7 cysteines which are responsible for forming disulfide bonds to create a homodimeric structure (19). Characterization of sequence/structure-function relationships of individual orthologs of H-IPSE is the subject of continued investigation.

Referred hyperalgesia is a unique feature of visceral pain which causes normally non-painful stimuli to feel painful, even in anatomically distant locations. Cyclophosphamide/ifosfamide administration in rodents is a well-established model of referred hyperalgesia (3–5). We have previously reported that intravenous delivery of H-IPSE^H06^ alleviates visceral and referred pain in ifosfamide-treated mice (17). In a similar fashion, bladder wall-injected H-IPSE^H03^ alleviated referred hyperalgesia in an IL-4 and NLS-dependent fashion. Post-operative pain did not affect the differences observed with H-IPSE^H03^ administered via bladder wall injection, as we were able to demonstrate a statistically significant increase in pain threshold (i.e., decreased referred hyperalgesia) in ifosfamide-treated mice who received H-IPSE^H03^. This suggests that bladder wall-injected H-IPSE^H03^ may have alleviated ifosfamide- and/or surgery-induced pain. Tail vein injection of H-IPSE^H03^ did not result in a significant difference between treatment groups (data not shown).

The voided spot on paper assay is a well-established, reliable model to assess lower urinary tract function in mice (1, 8, 11, 25). The characteristic voiding patterns of C57BL/6 mice consist of large volume voids in the corners of cages, whereas bladder injury causes mice to void at non-corner edges or the center of cages (25). We demonstrated that ifosfamide exposure alters voiding behavior by significantly decreasing corner voiding. This was non-significantly reversed by H-IPSE^H03^ bladder wall injection. Ifosfamide-treated mice tended to void in the central part of the cage, although this was not significant. The voided spot on paper assay results may have been influenced by surgical intervention and accompanying post-operative pain. Tail vein injection of H-IPSE^H03^ did not affect voiding patterns in ifosfamide-treated mice at all (data not shown). We did not allow mice to drink water during the 4 hour duration of assay to avoid possible interference of dripping water with collection of urine on filter paper. Post-operative pain in bladder wall-injected mice, as well as the lack of water access, may have influenced voiding behavior independent of the effects of IPSE.

We previously demonstrated that H-IPSE^H06^ induces transcription of uroplakins in the ifosfamide-injured bladder to a degree similar to or greater than MESNA (17). Uroplakins are transmembrane proteins implicated in barrier functions, urothelial proliferation and bladder regeneration (12). Co-incubation of a variety of urothelial cell lines with H-IPSE^H06^ significantly increases cell proliferation (data not shown). Moreover, co-incubation of the urothelial cell line MB49 with H-IPSE^H03^ induced a much stronger proliferative response than H-IPSE^H06^. The pro-proliferative effect of H-IPSE^H03^ was nuclear localization sequence-dependent. This supports the notion that H-IPSE^H03^ may upregulate urothelial repair mechanisms through translocation to the nucleus and modulation of gene expression. Future work will be directed towards further quantifying changes in uroplakin and related gene expression induced by H-IPSE^H03^.

This study has several limitations. For instance, the mechanism by which IPSE targets and sequesters chemokines is poorly understood. It is possible that IPSE’s chemokine-binding properties may play a role in its therapeutic effects in the ifosfamide-injured bladder. Furthermore, we have not elucidated the mechanism by which IPSE exerts its effects outside the bladder. We chiefly included experiments in which IPSE was delivered directly to the tissue of interest. It is unclear how the mechanism of action of H-IPSE^H03^ is different when administered in a local versus systemic fashion. For example, basophil and mast cell recruitment to the bladder to release anti-inflammatory IL-4 may be modulated differently based on the concentration of regional and systemic IPSE. Nuclear translocation is an important component of the therapeutic effect of IPSE and we have not yet investigated transcriptional regulation by H-IPSE^H03^. It is also unclear whether IPSE operates on a transcriptional level or whether there is a post-translational component to its mechanism of action. These are areas of ongoing work.

In summary, we report the potential therapeutic application of a parasite-derived protein, H-IPSE^H03^, to treat hemorrhagic cystitis via bladder wall injection. Because we have found that IPSE modulates the host immune system to dampen inflammation as well as pain responses, we speculate that this protein could potentially be used for broader indications, such as interstitial cystitis and other bladder pain syndromes.

## Acknowledgements

We thank H. Gil Rushton and Hans G. Pohl for their support of urologic basic science research at Children’s National Medical Center.

## Abbrevations

CFSE: 5-(and 6)-Carboxyfluorescein diacetate succinimidyl ester
IPSE: Interleukin-4 inducing principle from Schistosoma mansoni eggs
MESNA: 2-mercaptoethanesulfonic acid
NLSNuclear: localization sequence

